# Canine transmissible venereal tumor genome reveals ancient introgression from coyotes to arctic sled dogs

**DOI:** 10.1101/350512

**Authors:** Xuan Wang, Bo-Wen Zhou, Ting-Ting Yin, Fang-Liang Chen, Ali Esmailizadeh, Melissa M. Turner, Andrei D. Poyarkov, Peter Savolainen, Guo-Dong Wang, Qiaomei Fu, Ya-Ping Zhang

## Abstract

Dear Editor,

Ancient genome-sequencing studies have extensively refined our understanding of the genetic histories and adaptive evolution of humans^1^, as well as the domestic process of livestock and crops^2^. Recent studies about ancient and modern canids all indicate that canids have intricate histories, and lots of canine populations disappeared during evolution^3–6^. Thus, ancient genomes from canine fossils are in high demand to clarify canine evolution and the processes leading to dog domestication. Canine transmissible venereal tumor (CTVT), the oldest known somatic cell line, is a living fossil, originating from cancer cells transmitted from a host to other canids during the mating process^7^. Clonal origin analyses hints that the original dog infected with CTVT (CTVT founder) came from an ancient sled dog or wolf population^7–9^. However, the genetic composition of the CTVT founder is still not clear.

In order to explore this issue, we applied whole genome sequencing (WGS) to two CTVT samples, their corresponding hosts, and 24 additional canids (Supplementary Note). Combined with published WGS data of two CTVT samples and high quality canine WGS data, we constructed a data set containing WGS data of four CTVT samples a 169-individual reference panel composed of worldwide gray wolves (*Canis lupus*), dogs (*Canis lupus familiaris*), coyotes (*Canis latrans*) and golden jackals (*Canis aureus*) (Supplementary Note, Table S1).

We firstly developed a pipeline *transmissible tumor genotyper* to obtain the per-site genotype for each tumor sample (Supplementary Note). All sites with the same genotype among four CTVT samples were used for ascertainment. Next, we successfully ascertained 18.9M biallelic single nucleotide polymorphisms (SNPs) for the CTVT common genotype using 24.5M SNPs from the reference panel (Supplementary Note). We denote these data Set A. SNPs from the CTVT data in set A could be treated as the founder’s germline polymorphisms because they showed a typical germline mutation signature (Supplementary Note, Figure S5).

We utilized population phylogeny analysis, principal component analysis (PCA) and outgroup *f3* (X, CTVT founder; Outgroup) statistical analysis to assess the genetic relationship between the CTVT founder and modern samples in set A (Supplementary Note). Our results reveal that the CTVT founder was more closely related to present-day arctic sled dogs than to any other populations (Figure S6-8), in accordance with very recent results^7, 8^. However, ADMIXTURE analysis showed that the CTVT founder also possessed an ancestral component found predominantly in non-dog populations, a result that we do not observe for any arctic sled dog (Supplementary Note, Figure 1A). Moreover, the CTVT founder did not cluster tightly with arctic sled dogs in the PCA analysis (Figure S7). These results imply that the CTVT founder belonged to a previously unknown arctic dog population that is not represented in the reference panel.

**Figure 1.**
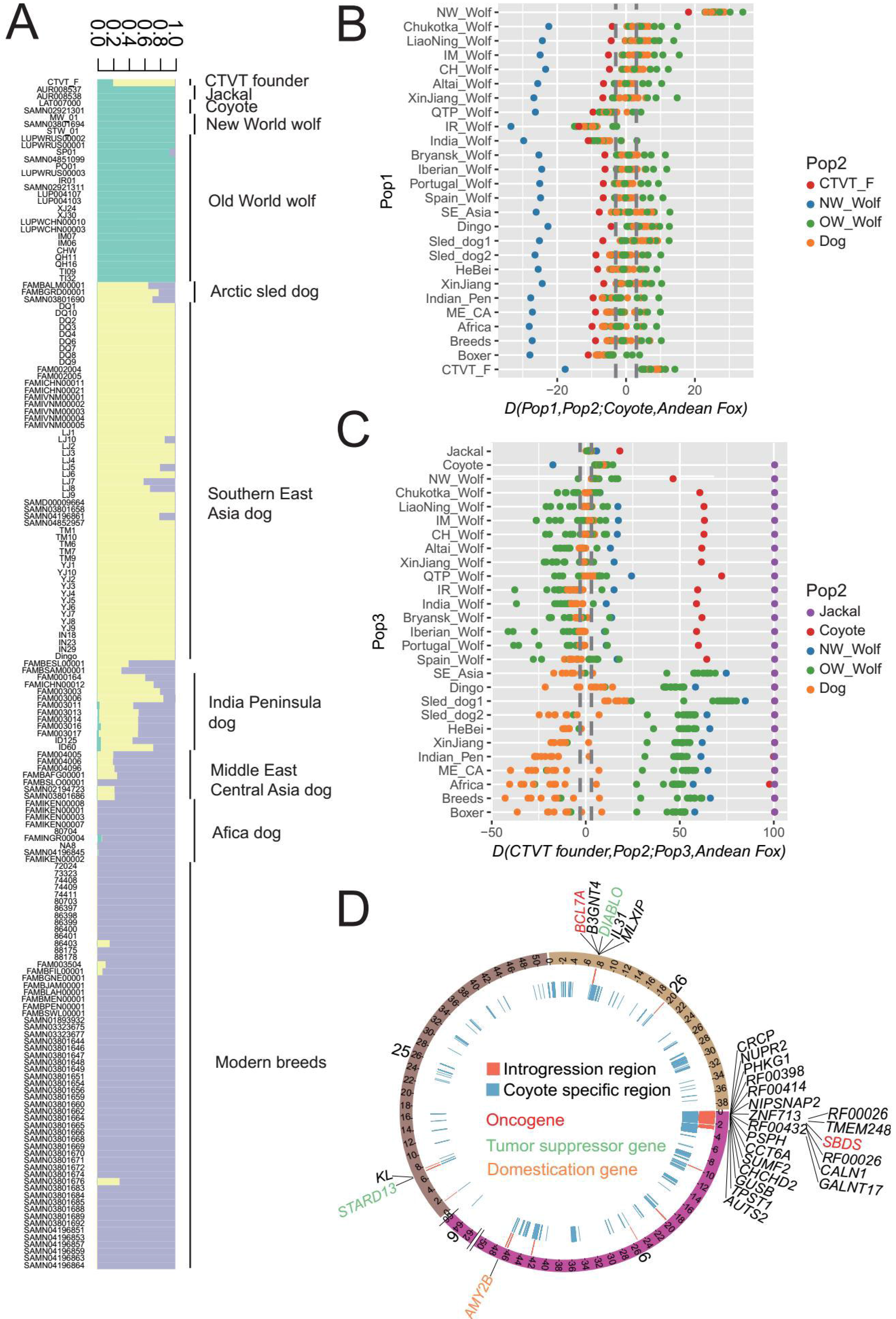
Analyses of introgression into the CTVT founder. **(A)** ADMIXTURE analysis result (K=3) for jackal, coyote, gray wolf and dog populations with 163K SNPs pruned by linkage disequilibrium. Colors indicate different ancestral components. **(B)** *D*(Pop1, Pop2; Coyote, Andean Fox), with the Z-score given on the x axis. When the CTVT founder is Pop2, Z>3 for all Pop1 groups except the New-World wolf which is a descendant of coyotes. **(C)** *D*(CTVT founder, Pop2; Pop3, Andean Fox), with the Z-score given on the x axis. When present-day arctic sled dogs and coyotes are Pop3, Z>3 scores for all Pop2 groups. **(D)** Introgressed segments (red regions) traced by high frequency coyote private allele positions (blue regions), see Supplementary Note. Genes within introgressed segments were labeled alongside ideogram of chromosomes. Copy number loss regions on chromosome 6 were truncated for clarity.

To further investigate whether the CTVT founder had introgression from a non-dog population, we ascertained set A using the Andean fox (*Lycalopex culpaeus*) genome (Supplementary Note), forming set B. Next, we tested every non-dog group in set B as candidate introgressor using *D* statistics of the form D(CTVT founder, Pop2; Introgressor, Andean Fox), where Pop2 was tested using all other present-day groups (Supplementary Note, Table S2). Only coyotes were found to be possible candidate introgressors, showing significant (Z>4.2) positive *D* statistics for all Pop2 populations except the New World wolves (Figure 1B). We then tested whether any domesticated dog populations could act as the candidate introgressor for the CTVT founder. Consistent with above results, we find that present-day arctic sled dogs showed significant (Z>9.6) positive D(CTVT founder, Pop2; Sled dog1, Andean Fox) statistics for all Pop2 populations (Figure 1C). *D* statistics suggest that the CTVT founder was an arctic sled dog bearing gene flow from a population most closely related to coyotes. We did not find strong evidence of coyote introgression into present-day arctic sled dogs (*D*(Sled dog1, Pop2; Coyote, Andean Fox) ~ 0). Finally, we used high frequency coyote private alleles to genotype the CTVT founder, in order to manually check whether there was any evidence of continuous coyote-related segments introgressed into the CTVT founder (Supplementary Note) and found one segment longer than 3Mbps on chromosome 6, and several shorter segments on chromosomes 6, 25 and 26. (Figure 1D). This finding helps us to reject the hypothesis that our results were caused by somatic reverse mutations, rather than introgression. 29 protein coding genes are located within these segments in total. Among them, we found four tumor-related genes^10, 11^ and a domestication gene^12^ *AMY2B* (Figure 1D).

The presence of coyote genotypes at tumor-related genes may contribute to the genetic susceptibility to CTVT tumorigenesis. The coyote genotypes at *AMY2B* may lead to low adaptation on a diet rich in starch and the disappearance of the population represented by the CTVT founder in agrarian age.

In conclusion, our detailed analyses reveal that the CTVT founder came from an arctic sled dog population that possessed introgression from a population related to coyotes, a result that was not known in previous studies. Considering the habitat of coyotes in North America^13^, we propose two hypotheses: 1) The CTVT founder lived in the arctic region of North America. 2) The CTVT founder lived in the arctic region of the Far East, where arctic dogs possessing the introgressed segments migrated through the Bering Strait in an unknown period. Hence, an ancient story of canine admixture is hidden in the genome of a living fossil, the CTVT. To further test our hypotheses of ancient admixture and to better understand the detailed evolutionary history of dogs from the arctic region and Americas, it is crucial to acquire ancient samples in these regions in future work.

## Acknowledgments

We thank Melinda A. Yang for her valuable comments, Newton O. Otecko for providing dog samples from Kenya, Ming-Shan Wang for collecting dog samples from Sri Lanka, Shi-Fang Wu for collecting other dog samples, Qi-Jun Zhou for collecting CTVT samples and laboratory work, and the BIG Data Center in Beijing Institute of Genomics, Chinese Academy of Sciences (CAS) for usage of their high-performance computing platform. This work was supported by the Strategic Priority Research Program (B) (XDB13000000) of the CAS, National Natural Science Foundation of China (91531303, 91731303, 41672021, and 41630102). Q.F. is supported by CAS (XDA19050102, QYZDB-SSW-DQC003, and XDPB05), and the Howard Hughes Medical Institute (grant number 55008731). G.D.W. is supported by the Youth Innovation Promotion Association, CAS.

